# Optimal Mass Transport Kinetic Modeling for Head and Neck DCE-MRI: Initial Analysis

**DOI:** 10.1101/612770

**Authors:** Rena Elkin, Saad Nadeem, Eve LoCastro, Ramesh Paudyal, Vaios Hatzoglou, Nancy Y. Lee, Amita Shukla-Dave, Joseph O. Deasy, Allen Tannenbaum

**Author notes:** Correspondence Saad Nadeem, Department of Medical, Physics, Memorial Sloan Kettering Cancer, Center, New York, NY 10065, USA. Equally contributing authors (Rena Elkin, Saad Nadeem). Funding information: AFOSR, Grant Number: FA9550-17-1-0435; National Institutes of Health, Grant Number: R01-AG048769 and R01-CA198121; MSK Cancer Center Support Grant, Grant Number: P30 CA008748; BRCF, Grant Number: BCRF-17-193.

## Abstract

Current state-of-the-art models for estimating the pharmacokinetic parameters do not account for intervoxel movement of the contrast agent (CA). We introduce an optimal mass transport (OMT) formulation that naturally handles intervoxel CA movement and distinguishes between advective and diffusive flows. Ten patients with head and neck squamous cell carcinoma (HNSCC) were enrolled in the study between June 2014 and October 2015 and underwent DCE MRI imaging prior to beginning treatment. The CA tissue concentration information was taken as the input in the data-driven OMT model. The OMT approach was tested on HNSCC DCE data that provides quantitative information for forward flux (Φ_*F*_) and backward flux (Φ_*B*_). OMT-derived Φ_*F*_ was compared with the volume transfer constant for CA, K^*trans*^, derived from the Extended Tofts Model (ETM). The OMT-derived flows showed a consistent jump in the CA diffusive behavior across the images in accordance with the known CA dynamics. The mean forward flux was 0.0082 ± 0.0091 (min^-1^) whereas the mean advective component was 0.0052±0.0086 (min^-1^) in the HNSCC patients. The diffusive percentages in forward and backward flux ranged from 8.67–18.76% and 12.76–30.36%, respectively. The OMT model accounts for intervoxel CA movement and results show that the forward flux (Φ_*F*_) is comparable with the ETM-derived K^*trans*^. This is a novel data-driven study based on optimal mass transport principles applied to patient DCE imaging to analyze CA flow in HNSCC.

## 1 INTRODUCTION

Head and neck (HN) tumors are heterogeneous with complex anatomy [1]. Accurate detection and delineation of tumor extent is critical to optimize treatment planning; patients therefore routinely undergo non-invasive imaging for careful assessment of this complex anatomy. Non-invasive magnetic resonance imaging (MRI) has served an important role as a diagnostic test for initial staging and follow-up tumors in the HN region [2, 3, 4].

Dynamic contrast-enhanced (DCE) MRI involves the acquisition of successive T1-weighted images with administration of a T1-shortening Gadolinium-based contrast agent (CA) [5]. DCE imaging has been considered to be a promising tool for clinical diagnostics, including head and neck cancers [6, 7]. DCE data can be evaluated by semi-quantitative analysis and model based quantitative analysis [5, 8]. Semi-quantitative analyses (ie, time to peak (TTP), maximum contrast enhancement) provide only limited information about tumor physiology [9].

The Tofts and Extended Tofts (ETM) pharmacokinetic models are the most commonly used models for quantitative DCE-MRI analysis [5]. In DCE-MRI, CA concentration is derived from changes in signal intensity over time, then fitted to a tracer model to estimate pharmacokinetic (PK) parameters related to tumor vascular permeability and tissue perfusion [5]. The volume transfer constant for CA (K^*trans*^) represents the contrast agent transport from the blood plasma into the extravascular extracellular space (EES), and has been shown to be a prognostic factor in head and neck cancer [6, 7].

The accuracy and precision of this quantitative parameter can be influenced by arterial input function (AIF) quantification, temporal resolution in data acquisition, signal-to-noise ratio (SNR), and model selection [10, 11, 12, 13, 14, 15, 16]. Moreover, these quantitative metrics are macroscopic, representing averages over highly heterogeneous microscopic processes, and requires an appropriate model to analyze the data.

The problem of optimal mass transport (OMT) was posed for finding the minimal transportation cost for moving a pile of soil from one location to another. OMT was given a modern and relaxed formulation with the Monge–Kantorovich problem [17, 18]. Applications include image processing and computer vision, econometrics, fluid flows, statistical physics, machine learning, expert systems, and meteorology [19, 20]. Benamou and Brenier [21] presented a dynamical version of OMT which has received much deserved attention. Their approach considers the minimization of kinetic energy subject to a continuity equation that enforces mass preservation.

It has been demonstrated that the differences in elevated tumor pressure lead to exudate convective flow out of the tumor such that any flow into the tumor may be ascribed to diffusion [22]. In the present work, we propose to employ methods from optimal mass transport (OMT) theory to study the problem of estimating CA flow from tumor DCE-MRI data. For the approach here, we add a diffusion term to the continuity equation. The resulting advection/diffusion equation regularizes the flow and moreover, as we will argue, gives a better physical picture of the underlying dynamics. This type of approach was exploited in our previous work [23] for the study of the glymphatic system.

We note that if one integrates the advection/diffusion equation over a given “compartment” and applies OMT theory, one derives an equation very similar to the ETM model [24]. In contrast to the conventional methods, the OMT framework provides dynamic information, formative of the time-varying directional nature of the flow. Other than using the advection/diffusion flow from computational fluid dynamics, we do not impose any prior constraints or assumptions of the underlying physics of the model; for example, we do not require AIF as an input. It is important to emphasize that our analysis is data driven, and all of our conclusions are based on the imagery.

## 2 METHODS

### 2.1 Optimal Mass Transport

The main tool we will be employing is based on the theory of optimal mass transport (OMT). We provide the theoretical details relevant to this work and simply refer the interested reader to [19, 20] for more details. A comprehensive list of symbols used in the following text and their interpretation is given in Table 1.

**TABLE 1.** Symbol Reference List

| Symbol | Interpretation |
| --- | --- |
| $C_t$ | tissue contrast agent concentration (mMol) |
| $C_p$ | plasma contrast agent concentration (mMol) |
| $v_e$ | volume fraction of extravascular extracellular space (EES) in Extended Tofts Model (ETM) |
| $v_p$ | volume fraction of vascular space in ETM |
| $K^{trans}$ | contrast agent volume transfer constant from the vascular space to the EES in ETM ($\text{min}^{-1}$) |
| $k_{ep}$ | contrast agent transport rate constant from the EES to the vascular space in ETM ($\text{min}^{-1}$) |
| $d$ | spatial dimension |
| $A$ | a subdomain of $\mathbb{R}^d$ |
| $x$ | spatial variable, $x \in A$ |
| $t$ | temporal variable |
| $\nabla \cdot (f)$ | spatial divergence of vector field $f : \mathbb{R}^d \rightarrow \mathbb{R}^d$ |
| $\nabla g$ | spatial gradient of scalar function $g : \mathbb{R}^d \rightarrow \mathbb{R}$ |
| $\lambda$ | potential function of the optimal velocity field |
| $D$ | symmetric positive definite diffusion matrix |
| $\sigma^2$ | diffusion coefficient ($\text{mm}^2/\text{min}$) |
| $\gamma$ | independent and identically distributed Gaussian noise with covariance $\Gamma$ |
| $\mu$ | noise free density |
| $\mu^\gamma$ | noisy density |
| $v$ | velocity |
| $\phi_F$ | forward flux ($\text{min}^{-1}$) |
| $\phi_B$ | backward flux ($\text{min}^{-1}$) |

For our purposes, we will review the fluid dynamical version of OMT for given initial and final densities associated with the same total mass, *µ*_0_ and *µ*_1_, defined on some domain *A* ⊂ ℝ^*d*^ over the normalized time interval *t* ∈ [0, 1] [21]:

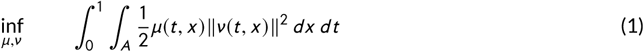

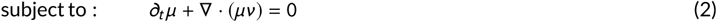

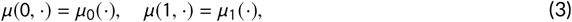

where *µ* = *µ*(*t*, *x*) ∈ ℝis the density, *ν* = *ν* (*t*, *x*) ∈ ℝ^*d*^ is the velocity, and the spatial dimension is taken to be *d* = 2 or *d* = 3. Here, the objective is to find the density *µ* and velocity *ν* characterizing the mass-preserving flow (2) with fixed temporal endpoints (3) that requires minimal expended kinetic energy (1). In simple terms, this approach seeks a flow over some time-space interval that minimizes the mass weighted kinetic energy of the flow and existence of the arguments *µ*_opt_ and *ν*_opt_ at which the infimum is obtained has been proven [21].

#### 2.1.1 Optimal Mass Transport with diffusion

We modify the original Benamou and Brenier OMT formulation by adding a diffusion term in the continuity equation (2). This allows for both advection and diffusion of the contrast agent and has the added benefit of regularizing the flow. Various aspects of regularized OMT have been studied in a number of recent works including [25, 26, 27, 28] which have extensive lists of references.

Accordingly, we substitute constraint (2) with the advection/diffusion equation

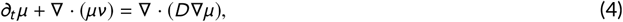

where *D*, encapsulating spatially-varying diffusion, is a positive definite symmetric matrix modelling the diffusion. We now formulate the ***regularized OMT*** problem as the infimum of the action given by (1) subject to the advection/diffusion constraint (4) with the given initial and final conditions (3).

#### 2.1.2 Hamilton-Jacobi equation

Using calculus of variations [20], one can show the optimal velocity for the regularized OMT problem is of the form

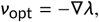

where *λ* satisfies the following diffused version of a Hamilton-Jacobi equation:

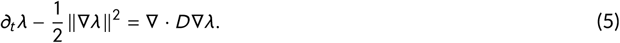

The latter equation derived from OMT (minimizing kinetic energy of the flow subject to an advection/diffusion constraint) is very revealing regarding the proposed model. Indeed, we note that the optimal velocity is the gradient of a function, and via (5), the bulk flow (advection) is clearly influenced as well by the diffusion. This will be described further in Section 2.1.3 below.

#### 2.1.3 OMT relation to Tracer Kinetic Model

We briefly sketch the relation to the very popular Tofts-Kermode model [5]. For simplicity, we indicate details in 2D. The basic idea is to integrate the advection/diffusion equation (4) over a given compartment *V* and use the divergence theorem. Accordingly, we let *S* denote the boundary of *V*. We first note that

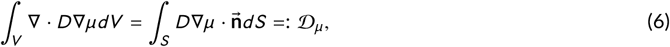

where 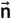 denotes the outward normal to the surface *S*.

**Remark:** In [29], 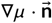 is interpreted as the difference between the concentrations of the contrast agent in the plasma and EES weighted by the ***permeability***, defined as K^*trans*^ [5], divided by the total surface area of the vasculature within the voxel multiplied by the volume of tissue within the voxel. This instantiation of Fick’s law serves to relate diffusive flux to differences in concentration.

Next we note that

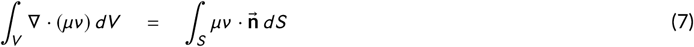

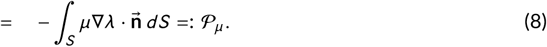

This part, derived from advection, may be interpreted as arising from Darcy’s law (difference in pressure) but because of (5), there is also the influence of diffusion. We should emphasize that OMT theory gives a unique solution of (4) with optimal velocity given as the gradient of a function via (5).

Finally, setting

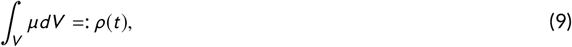

we see that at the optimal solution of the OMT problem, (4) becomes an ordinary differential equation of the form

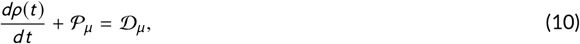

which leads to pharmacokinetic modelling.

Combining 𝒫_*µ*_ and 𝒟_*µ*_ (10), the rate of change of CA density over the compartment can be written as

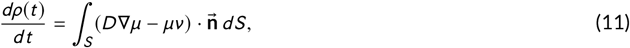

where 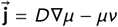 is the total flux. Although the direction of the flux, 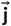, is influenced by both the density gradient ∇*µ* and the velocity *ν*, we chose to focus our attention on the velocity since it is affected by time-varying changes in the given images as well as diffusion, as seen from the Hamilton-Jacobi equation (5). Interestingly, we find that the velocity alone yields information that aligns well with the current understanding of the contrast behavior which we will elaborate on below.

### 2.2 Full OMT model for flow of contrast agent

In this section, we summarize our overall model based on the regularized OMT model. In what follows, we will take the diffusion matrix *D* = *σ*^2^*I*, that is, a constant times the identity. In other words, diffusion is not a tensor quantity, and there are no preferred directions.

In consideration of the data, we make the following assumptions to motivate the proposed regularized model:

i. A given sequence of *N* + 1 images taken at times *t* = *T*_*i*_ for *i* = 0, …, *N* are considered to be noisy observations of the CA’s true mass density at the respective times in which the images were captured:

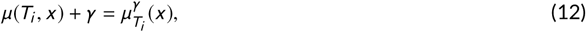

where *γ* is independent and identically distributed Gaussian noise with covariance 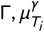 denotes the given noisy observed density and 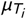 denotes the desired ‘true’ density.
ii. The flow is described by the advection/diffusion equation,

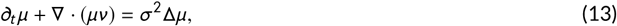

where *σ*^2^ ∈ ℝ is the diffusivity of the medium. We take *d* = 3 in what follows. Further, in our treatment above we took *T* = 1. Here we take a general final time *T*. The mathematical treatment is identical.

We propose to find the desired ‘true’ CA density interpolant *µ* : [0, *T*] × ℝ^*d*^ → ℝ between two given noisy observations 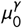 and 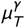 at times *t* = 0 and *t* = *T* respectively and the velocity field *ν* : [0, *T*] × ℝ^*d*^ → ℝ^*d*^ characterizing the evolution as the minimum arguments of the following action with a free endpoint term

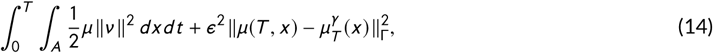

subject to the advection/diffusion constraint (13). The notation indicates that the variance operation is inversely scaled by the covariance. In this way, the low noise signals are more heavily weighted in the fitting to the observed values. The parameter *∊*^2^ is used to weight how harshly this data term should be penalized relative to the kinetic energy of the transport. We refer to (14) (with constraint (13) and fixed initial condition 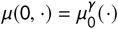) as the ***generalized regularized OMT*** problem (GR-OMT).

### 2.3 Patient Studies

The institutional review board approved and granted a waiver of informed consent for this retrospective clinical study, which was compliant with the Health Insurance Portability and Accountability Act. A total of 10 patients with neck nodal metastases and histologically-proven squamous cell carcinoma were enrolled into the study between June 2014 and October 2015. The 10 human papillomavirus positive (HPV+) patients were grouped as complete responders (CR) and non-CR based on the standard-of-care imaging and clinical follow-up according to the criteria in RECISTv1.1 [30].

### 2.4 DCE-MRI data acquisition

MRI studies were performed on a 3-Tesla (T) scanner (Ingenia; Philips Healthcare; Netherlands) using a neurovascular phased-array coil [31]. The standard MR acquisition parameters were as follows: multiplanar (axial, coronal, and sagittal) T2-weighted (T_2_w), fat-suppressed, fast spin-echo images (repetition time [TR]=4000 ms; echo time [TE]=80 ms; number of averages (NA)=2; matrix=256×256; slice thickness=5 mm; field of view [FOV]=20–24 cm), and multiplanar T1-weighted (T_1_w) images (TR=600 ms; TE=8 ms; NA=2; slice thickness=5 mm; matrix=256×256; slice thickness=5 mm; FOV=20–24 cm). Multiple flip angle pre-contrast T_1_w images were subsequently acquired for T_10_ measurement and DCE-MRI. Pre-contrast T_1_w images were acquired using a fast-multiphase spoiled gradient recalled (SPGR) echo sequence and the acquisition parameters were as follows: TR = 7 ms, TE = 2.7 ms, flip angles *θ* = 5°, 15°, 30°, single-excitation, NA = 1, pixel bandwidth = 250 Hz/pixel, 256 × 128 matrix that was zero filled to 256 × 256 during image reconstruction and FOV = 20-24 cm^2^.

The dynamic imaging sequence was acquired using the parameters for pre-contrast T_1_w with a flip angle of 15°. After acquisition of the first 5-6 images, antecubital vein catheters delivered a bolus of 0.1 mmol/kg Gd-DTPA (Magnevist; Berlex Labs, Montville, NJ) at 2 mL/s, followed by saline flush using an MR-compatible programmable power injector (Spectris; Medrad, Indianola, PA). Arterial concentration of contrast was not sufficient to induce an observable T_2_* effect; although this effect is present in high-field imaging (>3T), we assume that it is sufficiently small and exclude it from subsequent calculations. The entire node was covered with 5-mm thick slices, zero gap, resulting in collection of 8-10 slices with 7.46-8.1 second temporal resolution, for 40-60 time course data points. The time duration of dynamic imaging ranged 5-6 minutes.

### 2.5 DCE-MRI Theory

The T_1_-weighted (T_1_w) DCE signal for the spoiled gradient-echo recalled sequence is given by [32]:

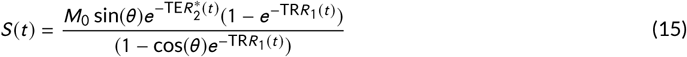

where *S* (*t*) is the voxel signal intensity at time *t*, *M*_0_ is the equilibrium magnetization of the protons, *θ* is the flip angle, TR is repetition time, and TE is echo time. *R*_1_(*t*) is the time course of longitudinal relaxation rate (*R*_1_ = 1/*T*_1_) and 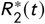 is time course of transverse relaxation rate 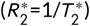. For 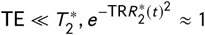.

The time course of measured signal is converted to the T_1_ relaxation rate. Under the fast exchange limit, the change in *R*_1_ (Δ*R*_1_ = *R*_1_ − *R*_10_) is linearly proportional to the tissue CA concentration *C*_*t*_ (mM).

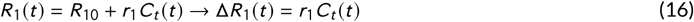

where *R*_10_ is the pre-contrast relaxation rate *R*_1_, and r_1_ is the longitudinal relaxivity of CA.

### 2.6 DCE-MRI Analysis

The *C*_*t*_ (*t*) was fitted to extended Tofts model (ETM) to estimate the physiological parameters. The *C*_*t*_ for ETM is given by [5]:

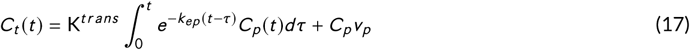

where *C*_*p*_ (*t*) is the time course of the plasma concentration of the CA (i.e., called arterial input function [AIF]), K^*trans*^ (min^-1^) is the volume transfer constant (vascular space to the EES), and *k*_*ep*_ = K^*trans*^ /*v*_*e*_ (min^-1^) is the rate constant for CA transport from the EES to vascular space. AIF for each patient was selected from the carotid artery using an automated cross-correlation routine to locate the single voxel with strongest bolus delivery signal (compared against a reference waveform) [33]. Regions of interest (ROI’s) on tumors were identified and contoured manually by an experienced neuroradiologist based on T_2_w images and late phases of the T_1_w DCE images. Tumor volume was calculated from the higher-resolution T_2_w images to lessen the effects of partial-volume on the estimate.

Pre-contrast T1 values (ie, T_10_) were calculated from the precontrast T1w variable flip angle images [34]. The time course of tissue concentration dataset, *C*_*t*_ (*t*), was fitted to Equation (17) using a nonlinear fitting technique that minimizes the sum of squared errors (SSE) between the model fit and data. Parameter estimation values were bounded in the fitting routine as follows: K^*trans*^ ∈ [0, 5] (min^-1^), *v*_*e*_, and *v*_*p*_ ∈ [0, 1]. The ETM parametric maps were generated using in-house MRI-QAMPER software (Quantitative Analysis Multi-Parametric Evaluation Routines) written in MATLAB.

### 2.7 OMT Numerical Simulation

We now outline our numerical implementation of the regularized OMT problem based on the work of [35].

We consider a cell-centered grid where cells are determined by voxels with volume *h*^*d*^. Linearized variables are denoted in bold and subscripts are used to indicate the time step (e.g. ***µ***_*n*_). As proposed in [35], we employ operator splitting to get the discretized advection/diffusion equation (D-ADE)

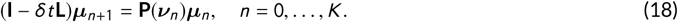

Here, *K* is the number of time steps, *δt* is the temporal discretization, **P** is a (tri)linear interpolation matrix that redistributes advected mass to the cell centers, **I** is the identity matrix and **L** is a discretized diffusion operator ∇ · *D* ∇. Even though we take the diffusion operator in the present paper to be of the form *σ*^2^Δ, we point out that the methodology works the same for a more general operator ∇ · *D* ∇ where *D* is a symmetric, positive definite matrix.

Given an initial density *µ*_0_ and velocity *ν*, the D-ADE (18) gives the density for any time step. Recalling that we fixed the initial density to be equal to the given image 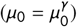, we see that the density can be written as a function of the velocity.

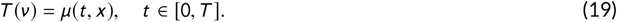

This allows us to solve the GR-OMT problem (14) as an unconstrained optimization problem for *ν* alone.

As shown in [35], the energy functional is discretized as

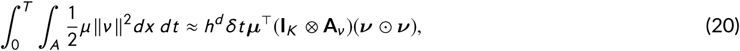

where ***µ*** and ***ν*** are block vectors comprised of the respective linearized variables ***µ***_*n*_ and ***ν*** _*n*-1_ for *n* = 1, …, *K*, **I**_*k*_ is the *k* × *k* identity matrix, **A**_*ν*_ is a 1 × *d* block matrix of **I**_*s*_, *s* is the number of voxels, ⊗ denotes the Kronecker product and ⊙ denotes the Hadamard product. Following [35], we solve the following discrete GR-OMT optimization problem:

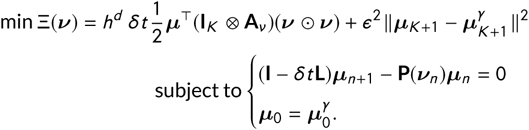

The constraint is linear with respect to ***ν*** and the objective function Ξ is quadratic with respect to ***ν*** and the interpolation matrix **P** is linear with respect to ***ν***. Thus one can employ a Gauss-Newton like method to solve the problem. Full details may be found in [35].

Recall that the GR-OMT algorithm yields the velocity characterizing the evolution between two given images, subject to the advection-diffusion equation (13). As described above, this velocity is influenced by both advective and diffusive behavior of the CA and looking at the average speed, (i.e. the magnitude of the velocity), turns out to be quite revealing. In particular, we define *forward flux* (Φ_*F*_, min^-1^) to be the average speed of the velocity at some initial time interval (CA delivery phase) and *backward flux* (Φ_*B*_, min^-1^) to be the average speed of the velocity for the remaining time interval; we investigated the selection of the specific time intervals for the most accurate assessment of Φ_*F*_ and Φ_*B*_. The flow diagram for the data processing steps in the GR-OMT pipeline is illustrated in Figure 1.

**FIGURE 1.**
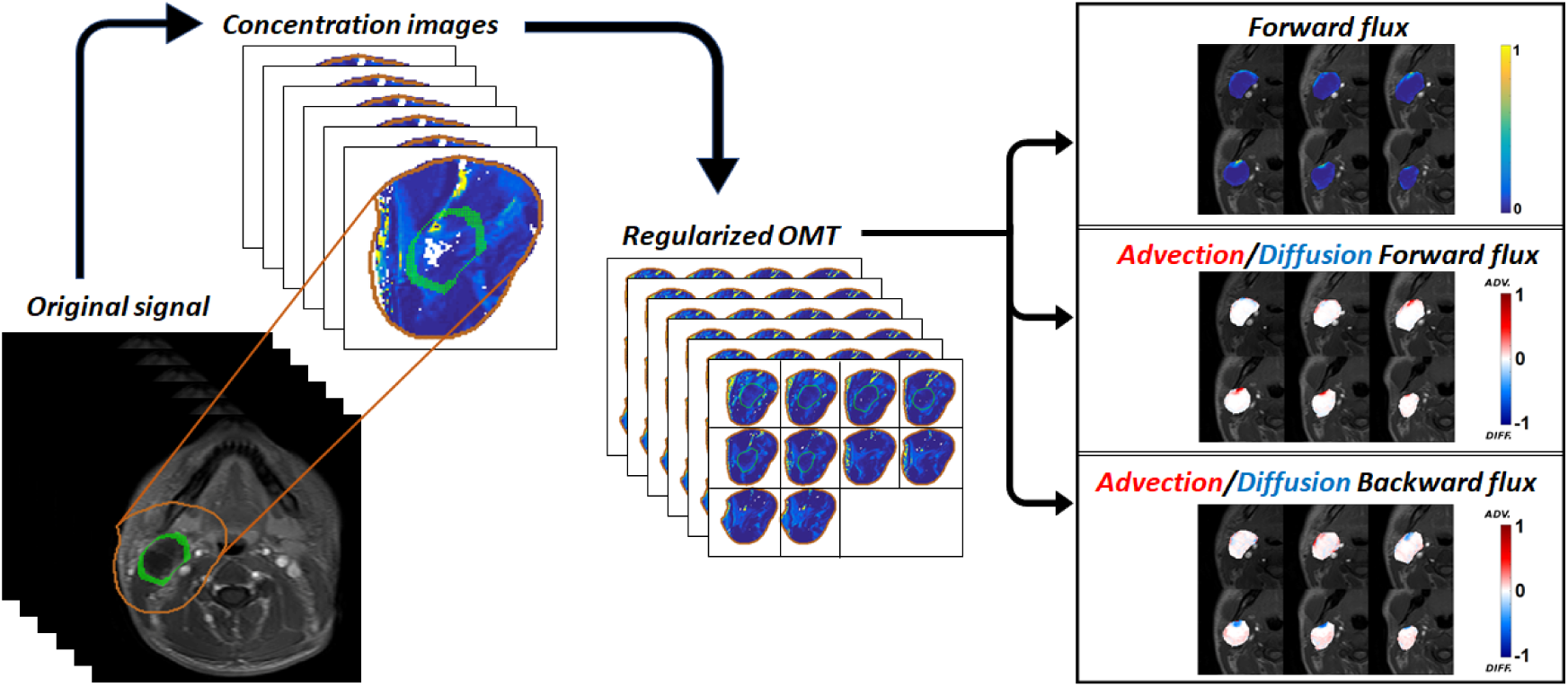
Optimal Transport pipeline. MR intensity images are converted to concentration images which are then fed to our regularized OMT algorithm. The resultant interpolations give ‘clean’ density maps and the corresponding velocity field is used to compute the forward flux by averaging the speed components over the delivery time points and pressure-driven advection/diffusion flows on the boundary via the directional component (advective flows are represented by movement out of the tumor whereas the diffusion is represented by movement into the tumor, i.e. against the pressure gradient).

A significant advantage of OMT framework is that we also get directional information from the velocity which cannot be inferred from imaging alone. In accordance with the finding that contrast is advectively driven out of the tumor and enters the tumor via diffusion due to differences in concentration gradient; a voxel is considered to be *advective* if the time-varying velocity vector at the corresponding location points out of the tumor for a majority of the time steps and *diffusive* if the velocity vector points inward for a majority of the time steps. The *diffusive percentage* is then defined to be the number of diffusive voxels divided by the tumor volume. Using smaller time intervals/increments in our pipeline, the diffusive percentage spike (indicating reflux) can be used to separate the backward and forward fluxes, as shown in Figure 2. Moreover, we had necrotic region segmentations for all 10 patient datasets and we observed that the speed via our analysis was zero in necrotic regions (meaning constant pressure) and non-zero in non-necrotic regions.

**FIGURE 2.**
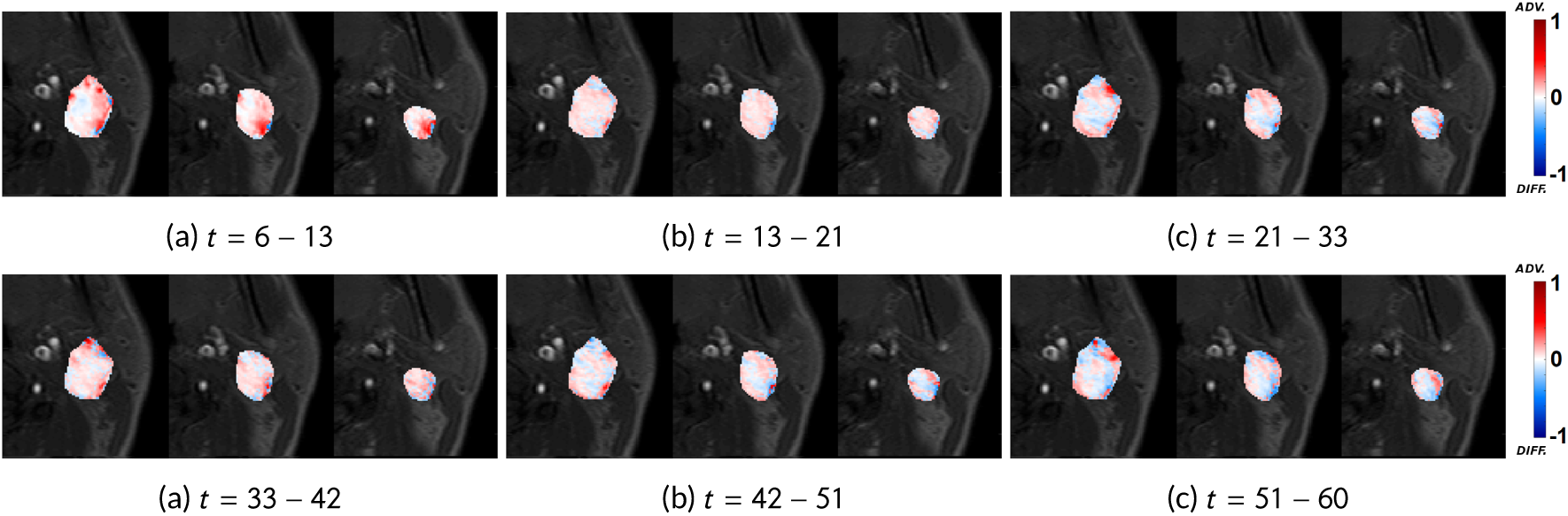
Smaller time increments. The diffusion percentages are (a) 18.76%, (b) 20.28%, (c) 25.14%, (d) 22.10%, (e) 23.66% and (f) 23.49%. We use the diffusion spike at (c) to divide the total time interval into *t* = 6 − 21 for forward flux and *t* = 21 − 60 for backward flux.

### 2.8 Statistical Analysis

Pearson correlation was performed between mean K^*trans*^ and mean Φ_*F*_ (with and without diffusion) values across all the 10 patients. The mean K^*trans*^ and mean Φ_*F*_ (with diffusion) values were strongly correlated (*r* = 0.79, p-value = 0.006). The mean K^*trans*^ and mean Φ_*F*_ (without diffusion) values were also found to be strongly correlated (*r* = 0.76, p-value = 0.01).

## 3 RESULTS

The mean volume of all tumors used in the study was 24.57 cm^3^ with standard deviation 9.51 cm^3^. For DCE-MRI images, we refer to the *i* -th image as the image taken at phase *i*. We selected image phases of *t* = 6−21, which typically represent the first pass of CA time interval, for forward flux and *t* = 21 − 39 for backward flux based on the observation that the diffusive percentage spikes at *t* = 21 across all our patients when looking at smaller time intervals/increments. As shown in Figure 2, the spike indicates the strong reflux (characteristic of backward flux). Figure 1 shows the OMT pipeline run where the corresponding forward and backward fluxes and their respective advection/diffusion components are illustrated. The mean tumor forward flux in this case is 0.0098 and the diffusive percentage in the delivery phase (*t* = 6 − 21) is 14.21% and increases to 22.66% in the remaining interval.

Figure 3 shows comparison of ETM-derived K^*trans*^ and OMT-derived Φ_*F*_. Abrupt changes in the neighboring slices/voxels are observed for K^*trans*^ since it ignores the intervoxel movement. In contrast, the Φ_*F*_ shows clearer signal strength and maintains the integrity of the neighboring voxels. As observed, K^*trans*^ follows similar trends to Φ_*F*_.

**FIGURE 3.**
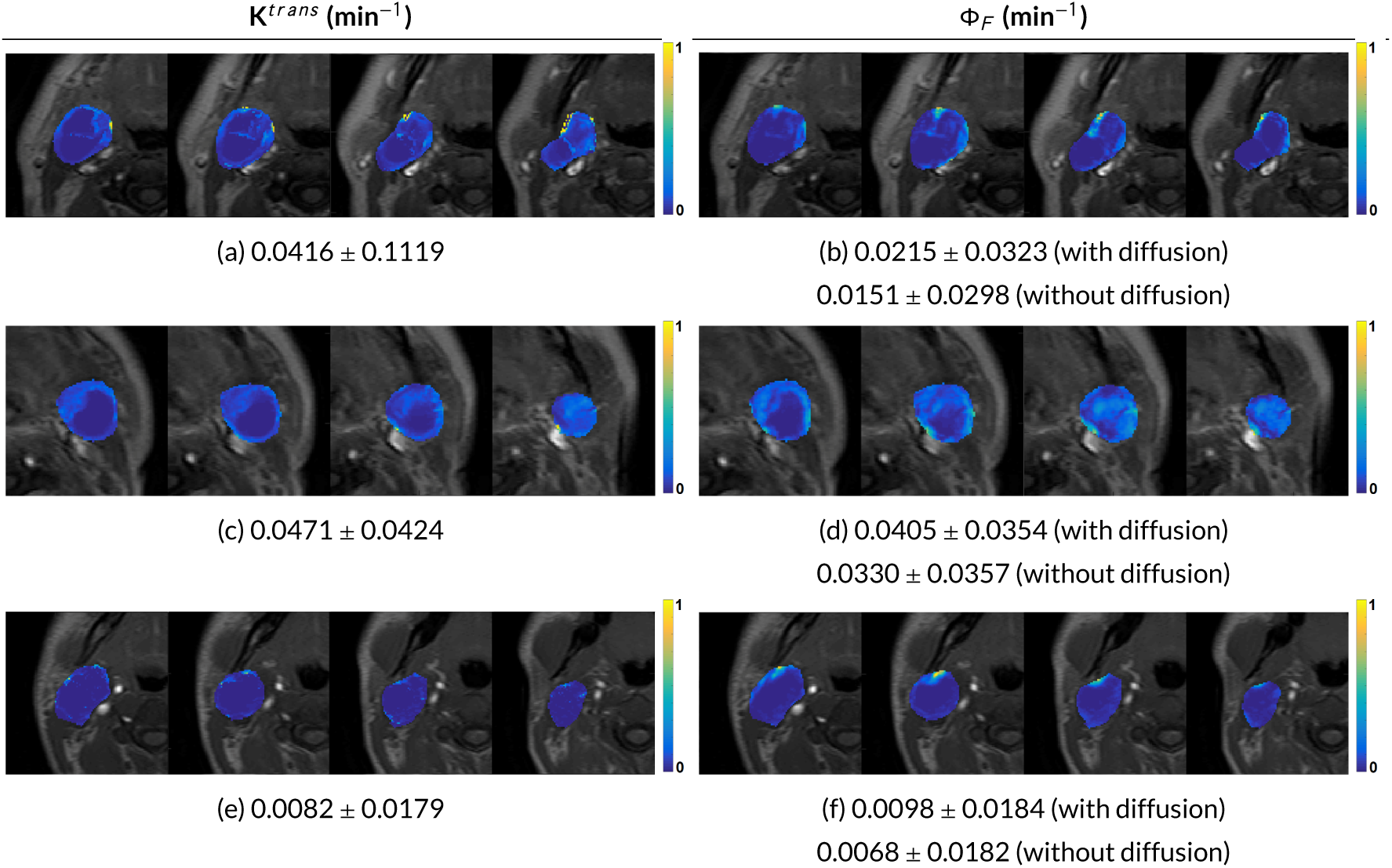
Comparison of mean±standard deviation for K^*trans*^ and forward flux (Φ_*F*_) values on 4 neighboring slices for 3 HNSCC patients. Note the abrupt changes in K^*trans*^ neighboring slices/voxels since it ignores the intervoxel movement. In contrast, forward flux shows clearer signals and maintains the integrity of the neighboring voxels.

We show forward and backward fluxes with corresponding advection and diffusion components for two patient datasets in Figures 4 and 5 based on the tumor size. Quantitative metrics for forward and backward flux for all the patients (used in this study) with and without diffusion are given in Table 2. The corresponding diffusive percentages for all the patients along with the tumor volumes and the CR/non-CR status are shown in Table 3. Out of 10 patients, there was only one patient (Patient ID: 8) who did not experience a complete response to the treatment, with the tumor detectable on six months standard follow-up imaging. This patient (Figure 5) exhibited the highest backward flux diffusive percentage, as shown in Table 3.

**TABLE 2.** Means and standard deviations of K^*trans*^, forward and backward fluxes for the 10 HNSCC patients. Mean forward/backward flux without diffusion = (sum of advective speeds)/(number of advective + diffusive voxels).

| | $K^{trans}$ | Forward Flux ($\Phi_F \text{ min}^{-1}$) | | Backward Flux ($\Phi_B \text{ min}^{-1}$) | |
| --- | --- | --- | --- | --- | --- |
| Patient ID | Mean ± Std. Dev. | with diffusion<br>Mean ± Std. Dev. | without diffusion<br>Mean ± Std. Dev. | with diffusion<br>Mean ± Std. Dev. | without diffusion<br>Mean ± Std. Dev. |
| 1 | 0.0324 ± 0.0348 | 0.0148 ± 0.0147 | 0.0099 ± 0.0139 | 0.0219 ± 0.0117 | 0.0102 ± 0.0134 |
| 2 | 0.0471 ± 0.0424 | 0.0405 ± 0.0354 | 0.0330 ± 0.0357 | 0.0311 ± 0.0279 | 0.0157 ± 0.0242 |
| 3 | 0.0047 ± 0.0052 | 0.0077 ± 0.0134 | 0.0063 ± 0.0118 | 0.0089 ± 0.0120 | 0.0061 ± 0.0113 |
| 4 | 0.0120 ± 0.0443 | 0.0221 ± 0.0267 | 0.0165 ± 0.0263 | 0.0176 ± 0.0288 | 0.0114 ± 0.0268 |
| 5 | 0.0416 ± 0.1119 | 0.0215 ± 0.0323 | 0.0151 ± 0.0298 | 0.0234 ± 0.0230 | 0.0131 ± 0.0214 |
| 6 | 0.0411 ± 0.0375 | 0.0470 ± 0.0412 | 0.0372 ± 0.0452 | 0.0438 ± 0.0410 | 0.0293 ± 0.0437 |
| 7 | 0.0225 ± 0.0211 | 0.0243 ± 0.0225 | 0.0167 ± 0.0223 | 0.0177 ± 0.0244 | 0.0089 ± 0.0215 |
| 8 | 0.0131 ± 0.0187 | 0.0082 ± 0.0091 | 0.0052 ± 0.0086 | 0.0103 ± 0.0108 | 0.0033 ± 0.0071 |
| 9 | 0.0082 ± 0.0179 | 0.0098 ± 0.0184 | 0.0068 ± 0.0182 | 0.0032 ± 0.0040 | 0.0014 ± 0.0026 |
| 10 | 0.0057 ± 0.0103 | 0.0107 ± 0.0151 | 0.0079 ± 0.0146 | 0.0124 ± 0.0151 | 0.0074 ± 0.0146 |

**TABLE 3.** Diffusive percentages (%) in forward and backward flux for the HNSCC patients with tumor volumes and complete responder (CR)/non-CR status.

| Patient ID | Tumor Volume (cm<sup>3</sup>) | Forward Flux ($\Phi_F$) | Backward Flux ($\Phi_B$) | CR Status |
| --- | --- | --- | --- | --- |
| 1 | 17.04 | 18.76% | 27.61% | CR |
| 2 | 25.23 | 8.67% | 17.42% | CR |
| 3 | 21.76 | 9.77% | 14.70% | CR |
| 4 | 28.28 | 9.96% | 20.16% | CR |
| 5 | 41.55 | 11.38% | 15.63% | CR |
| 6 | 55.27 | 11.83% | 16.60% | CR |
| 7 | 21.99 | 14.21% | 23.11% | CR |
| 8 | 15.04 | 15.20% | 30.36% | non-CR |
| 9 | 26.34 | 14.21% | 22.66% | CR |
| 10 | 66.63 | 10.78% | 12.76% | CR |

**FIGURE 4.**
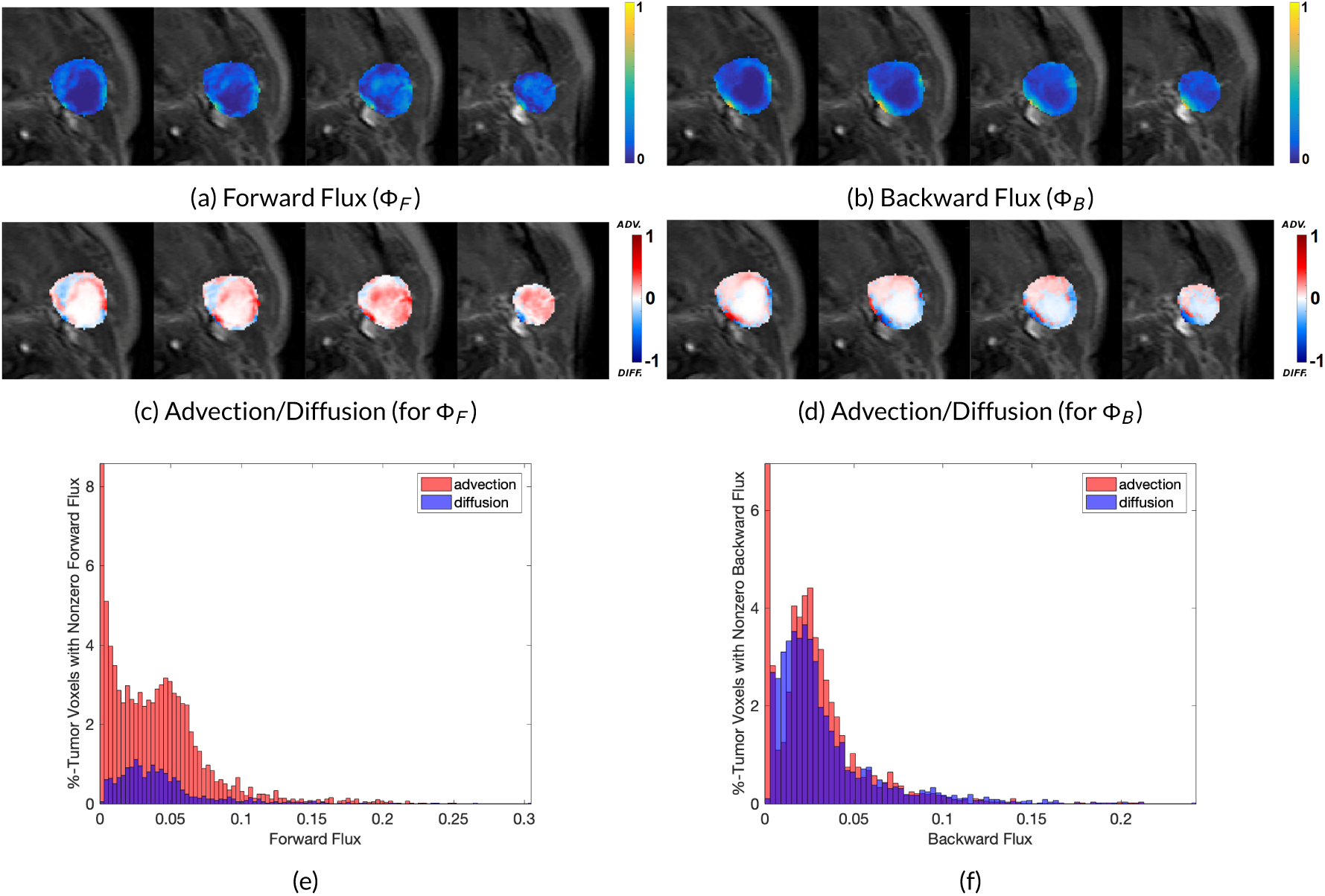
Forward and backward flux and the corresponding advection and diffusion components. The diffusion percentage increases from 8.67% in forward flux to 17.42% in backward flux. **(e)** Mean advective (red) and diffusive (blue) speeds given as the percentage of tumor voxels with nonzero mean speed (Forward Flux).

**FIGURE 5.**
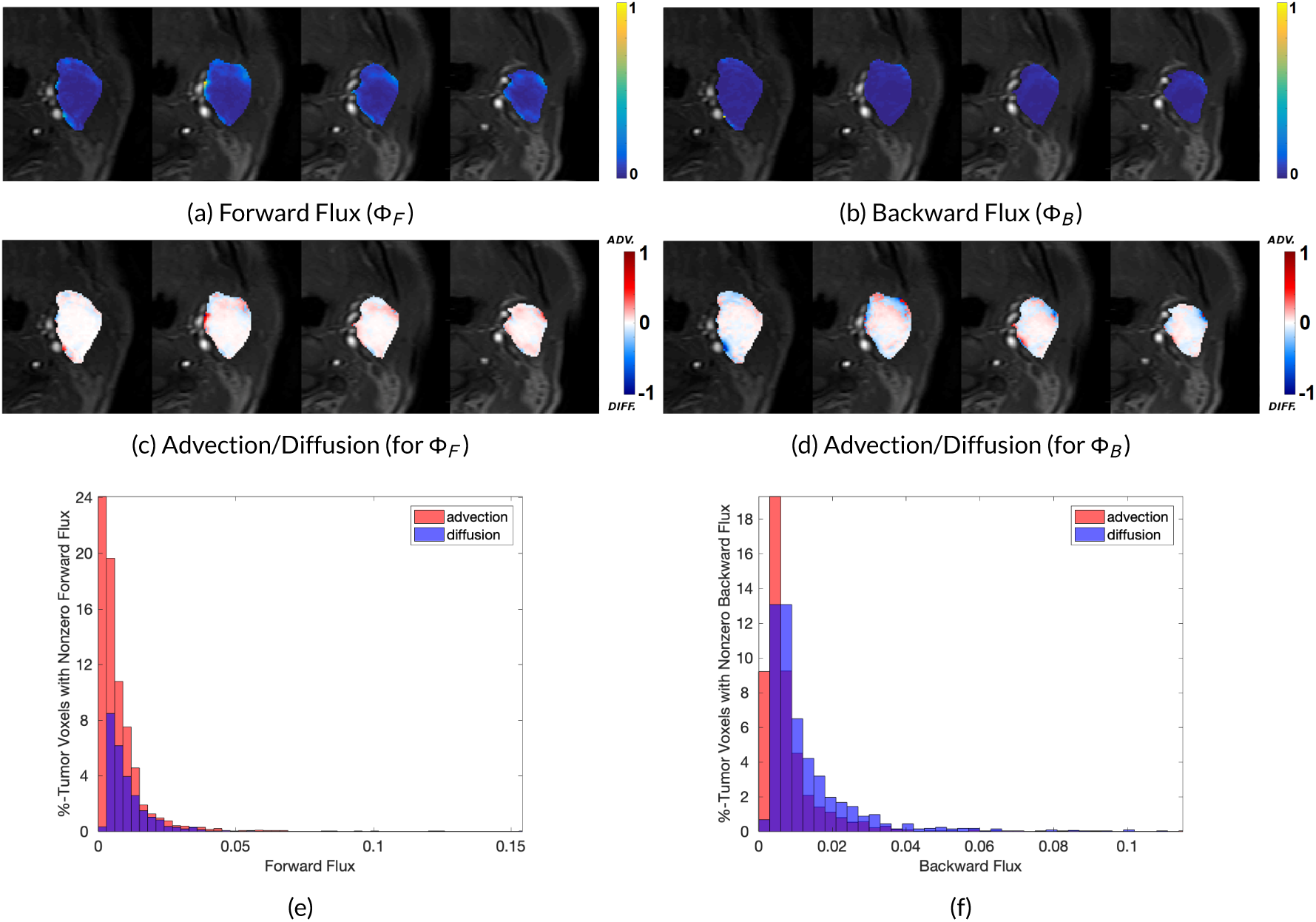
Forward and backward flux and the corresponding advection and diffusion components. The diffusion percentage increases from 15.20% in forward flux to 30.36% in backward flux. **(e)** Mean advective (red) and diffusive (blue) speeds given as the percentage of tumor voxels with nonzero mean speed (Forward Flux).

Finally, the sensitivity analysis of different values of *σ*^2^ is given in Figure 6. We see that the direction of the flux changes gradually as *σ*^2^ increases but remains stable. The movement becomes more uniform as the value increases with a large difference observed when *σ*^2^ = 1, as illustrated in (f). Given the stability of the low *σ*^2^ values, we used *σ*^2^ = 0.0001 for all the HNSCC datasets in this study. The *σ*^2^ = 0.0001 is half that of the diffusivity for gadolinium measured in an in-vivo preclinical study [36].

**FIGURE 6.**
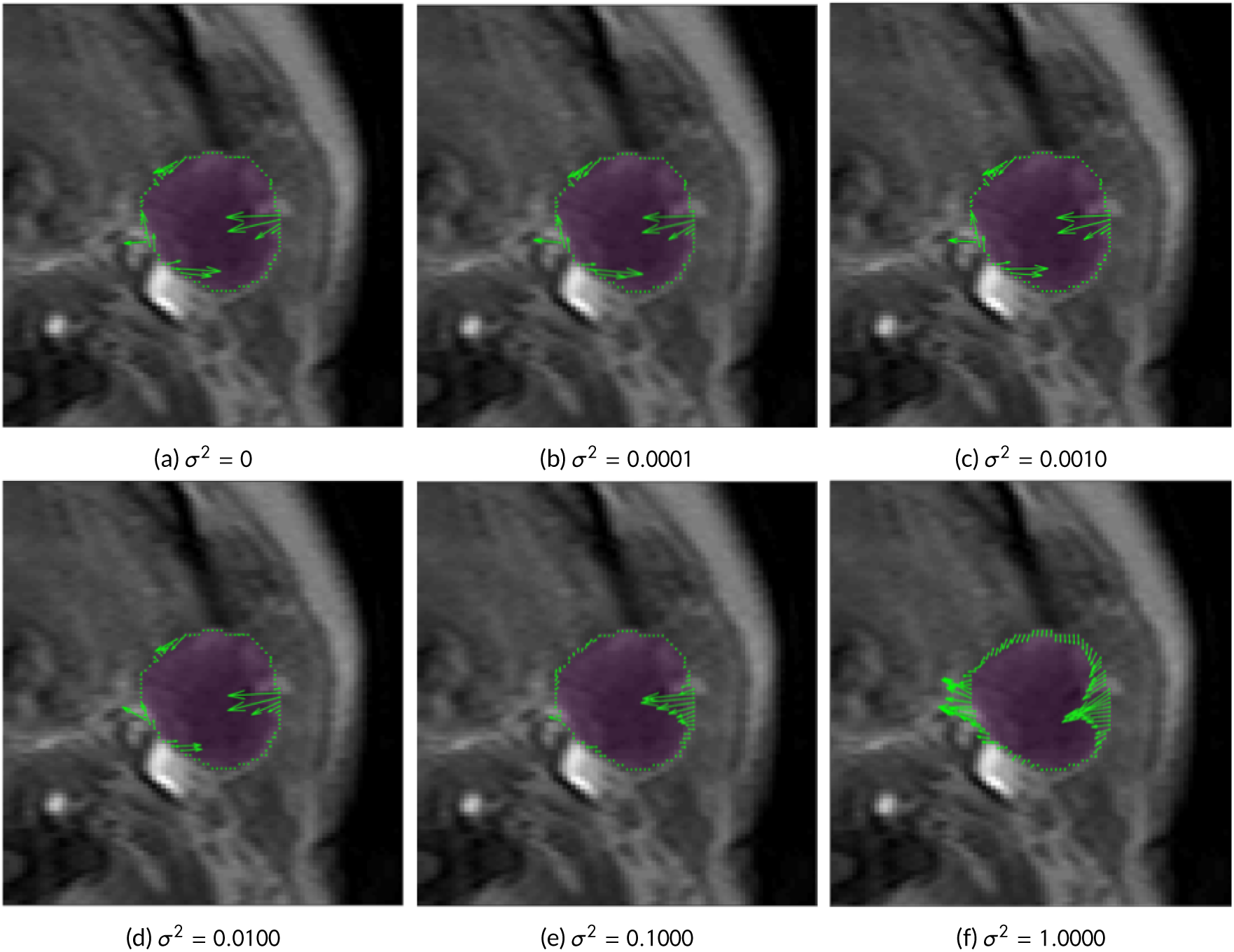
Comparison of flux *j* = *σ*^2^∇*µ* − *µν* for increasing values of *σ*^2^. The flux vectors are shown in green on the tumor boundary, highlighted in purple. We see that the direction of the flux changes gradually as *σ*^2^ increases but remains stable. The movement becomes more uniform as the value increases with a large difference observed when *σ*^2^ = 1, as illustrated in (f). Given the stability of the low *σ*^2^ values, we used *σ*^2^ = 0.0001 for all the HNSCC datasets in this paper.

## 4 DISCUSSION

In this paper, we have described a natural fluid flow model applied to DCE-MRI head and neck data. The model, unlike ETM, allows to take into account both advective and diffusive flows between neighboring voxels. In our framework, we directly applied an advection/diffusion model to study the optical flow properties of the dynamic imagery with no extraneous parameters. Thus the methodology described in the present work is data driven. We have shown how the bulk motion is chained to the diffusion through the diffused Hamilton-Jacobi equation (5). The relation to the compartmentalized models are revealed in (11) that shows explicitly how the time course of CA concentration is effected by a general velocity term that captures the conservation principles underlying the compartmental models.

ETM takes an input, AIF, a signal curve timecourse of CA delivery from a pre-selected arterial region and uses this as convolution for all the voxels in the tumor ROI. We do not make any such assumption. AIF for ETM and our analysis gave the same timeframe for initial pass of CA delivery in all the HNSCC datasets. ETM provides estimate of K^*trans*^ based on the non-linear fitting. The reliance on initial value estimates (AIF and T1 maps) in nonlinear least squares fitting is a major difference from the data driven OMT model employed in this study.

There is another way of looking at our approach, which may illuminate some of the key ideas. Optimal mass transport defines a metric on distributions that allows natural physically based interpolations of data. Mathematically, it defines a geodesic path in the space of distributions in a certain precise sense [37]. However, it does not take into account diffusion. Adding diffusion to the model not only regularizes the OMT framework, but adds another important interpretation to our proposed method. Namely, (4) may be regarded as a Fokker-Planck equation, and hence describes the transition probabilities of the associated stochastic differential equation [38]. Thus, instead of moving along geodesic paths, we are now moving along Brownian paths [25]. Therefore, the model we employ seems quite natural, since our data are noisy, and we do not want to impose any a priori constraints. Of course, data driven approaches, under which our proposed method falls, have become very popular in the treatment of many problems in engineering and applied science.

In the future, we will validate our measures using preclinical data and do a multiparameteric comparison (including PET/CT). We also plan to estimate the pressure directly from the presented approach. Moreover, we will add source terms to our formulation that will allow us to combine OMT with other metrics such as *L*^2^ and Fisher-Rao [39] while still keeping the underlying advection/diffusion model governing the flow. Finally, we will consider more general versions of the diffusion coefficient, e.g. taking into account spatial dependence in order to further improve our scheme for the kinetic modeling of tumor DCE data.

## ACKNOWLEDGEMENTS

This project was supported by AFOSR grant (FA9550-17-1-0435), grants from National Institutes of Health (R01-AG048769, R01-CA198121), MSK Cancer Center Support Grant/Core Grant (P30 CA008748), and a grant from Breast Cancer Research Foundation (grant BCRF-17-193).

## CONFLIC TO FINTEREST

There are no conflicts of interest.

